# Biofabricating murine and human myo-substitutes for rapid volumetric muscle loss restoration

**DOI:** 10.1101/2020.05.25.114819

**Authors:** Marco Costantini, Stefano Testa, Ersilia Fornetti, Claudia Fuoco, Minghao Nie, Sergio Bernardini, Alberto Rainer, Jacopo Baldi, Carmine Zoccali, Roberto Biagini, Luisa Castagnoli, Libero Vitiello, Bert Blaauw, Dror Seliktar, Wojciech Święszkowski, Piotr Garstecki, Shoji Takeuchi, Gianni Cesareni, Stefano Cannata, Cesare Gargioli

**Affiliations:** Institute of Physical Chemistry, Polish Academy of Sciences, Warsaw, Poland; Department of Biology, Rome University Tor Vergata, Rome, Italy; Department of Mechano-Informatics, Graduate School of Information Science and Technology, The University of Tokyo, Tokyo, Japan; Faculty of Engineering, Università Campus Bio-Medico di Roma, Rome, Italy; IRCCS Regina Elena National Cancer Institute, Rome, Italy; Department of Biology, University of Padova, Padova, Italy; Department of Biomedical Science and Venetian Institute of Molecular Medicine, University of Padova, Padova, Italy; Department of Biomedical Engineering, Techion Institute, Haifa, Israel; Faculty of Materials Science and Engineering, Warsaw University of Technology, Warsaw, Poland; Institute of Industrial Science, The University of Tokyo, Tokyo, Japan; IRCCS Fondazione Santa Lucia

**Author notes:** These authors have contributed equally (Marco Costantini, Stefano Testa).

## Abstract

The importance of skeletal muscle tissue is undoubted being the controller of several vital functions including respiration and all voluntary locomotion activities. However, its regenerative capability is limited and significant tissue loss often leads to a chronic pathologic condition known as volumetric muscle loss. Here, we propose a biofabrication approach to rapidly restore skeletal muscle mass, 3D histoarchitecture and functionality. By recapitulating muscle anisotropic organization at the microscale level, we demonstrate to efficiently guide cell differentiation and myobundle formation both *in vitro* and *in vivo*. Of note, upon implantation, the biofabricated myo-substitutes support the formation of new blood vessels and neuromuscular junctions – pivotal aspects for cell survival and muscle contractile functionalities – together with an advanced along with muscle mass and force recovery. Together, these data represent a solid base for further testing the myo-substitutes in large animal size and a promising platform to be eventually translated into clinical scenarios.

## 1. Introduction

Skeletal muscle (SM) tissue accounts for over 40% of the total body mass, controlling all voluntary locomotion activities and regulating vital functions such as respiration and metabolic homeostasis. Its functionality is therefore essential to maintain an adequate life quality [1–4].

The clinical scenarios of SM-related disorders are complex as the functionality of this tissue can be impaired by several conditions that can either be acquired, as a consequence of contusions, lacerations or surgical interventions (sarcoma removal), or be congenital/genetic as in muscular dystrophies or metabolic diseases. Regardless the disorder origin, treating any type of SM loss or disorder remains an unmet therapeutic need [5,6].

In this context, skeletal muscle tissue engineering (often referred as SMTE) holds great promises for addressing the two key challenges in the field: i) the development of a reliable 3D *in vitro* models and ii) the biofabrication of a macroscopic engineered SM tissue (construct volume > 1 cm^3^) [7–9].

As far as the first challenge is concerned, SMTE is expected (likely) to yield in the near future functional *in vitro* models for developing and testing approaches that may help addressing muscle dysfunctions caused by genetic or metabolic diseases. The success in the development of dependable *in vitro* models of human SM, aside from contributing to sparing animal experimentation, it is expected to overcome some of the problems caused by the failure of some animal models and to reproduce some important features of the human disease. In addition, such *in vitro* models will therefore be of great help in better understanding the etiology of SM specific diseases – including dystrophies such as Sarcoglycanopathy or Facio-Scapulo-Humeral-Dystrophy and metabolic disorders such as Pompe disease and Mitochondrial Metabolic Myopathy [10–12]. Finally, in the context of drug discovery and development, SMTE-based *in vitro* models will be especially valuable for the characterization of drug molecular mechanisms, rapid prioritization of lead candidates, toxicity testing and new biomarker identification [13].

The development of *in vitro* SM tissue models has been at the core of intensive industrial and academic research over the past ten years and an ever-increasing number of innovative approaches have been proposed including 3D bioprinting and organ-on-a chip solutions [14–17].

Despite the introduction of new technologies and refined protocols for culturing engineered muscles – derived from either murine or human muscle progenitors – so far, SM maturation and maintenance have been only partially recapitulated *in vitro* and, most importantly, to a small scale [18,19]. These approaches, while promising, are still far from satisfactorily addressing the second key challenge of SMTE – i.e. the development of macroscopic tissue-equivalents of a size suitable for treating volumetric muscle loss (VML): a condition resulting from large traumatic injury or massive surgical ablation upon tumor removal [20,21]. VML, which is characterized by muscular mass loss, functional deficit and permanent disability, is currently treated by surgical tissue transfer (autologous transplantation). This invasive approach is often associated with unsatisfactory outcomes due to poor tissue engraftment and donor site morbidity [22–24].

In this study, we present a microfluidic wet-spinning system [25] that allows rapid fabrication (fiber extrusion velocity > 2 m/min) of macroscopic yarns of cell-laden hydrogel microfibers – loaded either with murine (mesoangioblasts, Mabs) or human (primary myoblasts) muscle precursors – that closely mimic the structure of the SM tissue. Our findings demonstrate that by optimizing key parameters such as i) hydrogel precursor composition (i.e. bioink), ii) cell seeding density, iii) fiber architecture and iv) culturing protocol, we are able to successfully fabricate advanced artificial myo-substitutes that can be used to restore SM mass-loss and functionality *in vivo*.

## 2. Material and Methods

### 2.1 Materials

All chemicals were purchased from Sigma-Aldrich and used without further purification unless otherwise stated. Sodium alginate (ALG, Mw 33 kDa) was a kind gift from FMC Biopolymers. Photocurable PEG-fibrinogen was synthesized following previously published protocols [26].

### 2.2 Design and fabrication of wet-spinning set-up

Cell-laden hydrogel fibers were fabricated using a custom wet-spinning approach [25]. The whole system is composed of a co-axial nozzle 3D printed via stereolithography (DWS, DIGITALWAX 028J, Italy using a THERMA DM210 photoresin) and a stepper motor control through an Arduino UNO electronic board. The characteristic dimensions of the co-axial nozzle were as follows: ID 0.3 mm and OD 1.3 mm. The internal needle protrudes out from the external one of approximately 500 μm [27]. After 3D printing, the nozzle was thoroughly rinsed in ethanol to remove unreacted photoresin and then dried at room temperature overnight. Prior to use, the co-axial nozzle was additionally coated with parylene (coating thickness ≈ 2 μm) using a chemical vapor deposition machine (Parylene Deposition System 2010, Specialty Coating Systems Inc.). Finally, the 3D printed needle was assembled on top of a polycarbonate structure and placed centrally with respect to the rotating Teflon drum connected to the stepper motor shaft.

### 2.3 Biofabrication of 3D constructs

#### Bioink formulation

All the experiments were performed using an alginate-based bioink (ALG = 4% w/v) blended with photocurable PEG-fibrinogen (PEG-Fib = 0.8% w/v). Biopolymers were dissolved in a 25 mM HEPES buffer solution and 0.1% w/v Irgacure 2959 was added to the bioink as radical photoinitiator. Skeletal muscle progenitors (human primary myoblasts or murine mesangioblasts) were resuspended in the bioink to a final concentration of 2 x 10^7^ cells/mL. Such cell density has been thoroughly optimized in our previous work [7].

#### Bulk hydrogel fabrication

3D bulk hydrogels were fabricated by a casting method. Briefly, bioink solution loaded with cells were poured into cylindrical silicon molds (PDMS) and then exposed first to a CaCl^2+^ solution (to promote physical alginate crosslinking) and then for 5 minutes to UV light (to polymerize PEG-fibrinogen).

#### 3D wet-spun fiber construct fabrication

Wet-spun hydrogel fiber scaffolds were fabricated using the custom set-up described in the paragraph 2.2. Prior to perform experiments with cells, the wet-spinning system was deeply characterized to investigate the relations between motor speed, fluid flowrates (bioink and calcium chloride solution) and fiber size. A typical experiment consists in supplying the bioink through the inner nozzle and a calcium chloride solution (0.3 M) through the external one. When the two solutions come into contact at the tip of the co-axial nozzle system, an instantaneous gelation of the bioink solution occurs. The resulting hydrogel blob is then pulled gently with a tweezer – forming a hydrogel fiber – until it reaches the surface of the rotating Teflon drum connected to the motor shaft. As soon as the fiber touches the drum, a hydrogel fiber starts to be continuously extruded from the nozzle and collected onto the drum forming a bundle. After careful optimization, we decided to fabricate hydrogel fibers of around 100 mm. Fluid flowrates and motor speed were adjusted as follows: bioink flowrate = 50 μL/min, calcium chloride flowrate = 25 μL/min and motor speed = 50 rpm. Samples were collected for approximately 2 min. After that, they were UV crosslinked to stabilize the PEG-fibrinogen.

### 2.4 In vitro cell culture

nLacZ modified mouse mesoangioblasts (Mabs) were obtained as already described in previous work (Fuoco et al., 2012). Briefly, Mabs were transduced by nLacZ lentiviral particles and cultured on Falcon dishes at 37°C with 5% CO2 in DMEM GlutaMAX (Gibco) supplemented with heat-inactivated 10% fetal bovine serum (FBS), 100 international units/mL penicillin and 100 mg/mL streptomycin. Human muscle derived primary myoblasts were isolated and culture as previously described[28]. Briefly, human muscle biopsies from healthy donors were minced and digested in 0.8% w/v collagenase I (Life Technologies; Carlsbad, CA, USA) in DMEM, supplemented with 100U/ml penicillin, 100mg/ml streptomycin (Life Technologies; Carlsbad, CA, USA), for 60 min. Afterwards, muscle fragments were gently dissociated by pipetting and passing through a 21 G syringe needle. Cell suspension was centrifuged for 10’ at 300g and the pellet was resuspended and seeded on a matrigel-coated 35 mm Petri dishes in growth medium composed of 20% FBS, 25ng/μl hFGFb (human Basic Fibroblast Growth Factor, Immunotools; Friesoythe, Germany) in Ham’s F12 medium (Euroclone; Milan, Italy) with Pen/Strep. Cells were expanded and cultured in 100 mm Petri dishes.

### 2.5 In vivo construct implantation

Two-month-old male SCID/Beige mice were anesthetized with an intramuscular injection of physiologic saline (10 ml/kg) containing ketamine (5 mg/ml) and xylazine (1 mg/ml) and then the 3D constructs were implanted in Tibialis Anterior muscle (TA), according to following surgical procedure [29]: a) limited incision on the medial side of the leg has been performed in order to reach the TA; b) utilizing a cautery to avoid bleeding, the muscle fibers were deeply removed to create a venue for the implant; c) the construct was placed in the removed TA fibers lodge and the incision was sutured. As control, the contralateral TA was surgically ablated but no construct was implanted. Analgesic treatment (Rimadyl, Pfizer, USA) was administered after the surgery to reduce pain and discomfort. Mice were sacrificed 20 days after implantation for molecular and morphological analysis. Experiments on animals were conducted according to the rules of good animal experimentation I.A.C.U.C. no 432 of 12 March 2006 and under Italian Health Ministry approval n° 228/2015-PR.

### 2.6 Force measurements

Contractile performance of treated and control muscles was measured *in vivo* using a 305B muscle lever system (Aurora Scientific, Aurora, ON, Canada) in animals anesthetized with a mixture of tiletamine, zolazepam, and xilazine. Mice were placed on a thermostatically controlled table, with knee kept stationery and foot firmly fixed to a footplate, which, in turn, was connected to the shaft of the motor. Contraction was elicited by electrical stimulation of the peroneal nerve. Teflon-covered stainless-steel electrodes were implanted near the branch of the peroneal nerve as it emerges distally from the popliteal fossa. The two thin electrodes were sewn on both sides of the nerve, and the skin above was sutured. The electrodes were connected to an AMP Master-8 stimulator (AMP Instruments, Jerusalem, Israel). Isometric contractions induced at different frequencies were performed to determine the force-frequency relationship.

### 2.7 Histology and immunostaining

*In vitro* 3D constructs after 20 days of culture, were fixed in PFA 2%, while *in vivo* grafts were processed for histology as previously described [7]. Briefly, 3D constructs were surgically explanted, embedded in O.C.T. and quickly frozen in liquid nitrogen cooled isopentane for sectioning at a thickness of 10 μm on a Leica cryostat. Then *in vitro* and *in vivo* resulting samples were processed for immunofluorescence. As primary antibodies, we used anti-Myosin Heavy Chain (mouse monoclonal, DHSB, diluted 1:2), anti-Laminin (rabbit polyclonal, Sigma-Aldrich # L9393, diluted 1:100), anti-Dystrophin Rod Domain (mouse monoclonal, Vector Laboratories # VP-D508, diluted 1:100), anti-von Willebrand factor (rabbit polyclonal, Dako # A0082, diluted 1:100), anti-Pax7 (mouse monoclonal, DHSB, diluted 1:4) and anti-Neurofilament H phospho (rabbit polyclonal, BioLegend # PRB-573C, diluted 1:400). As secondary antibodies, we used antimouse Alexa Fluor 555^®^ (goat polyclonal, Invitrogen # A32727, diluted 1.200) and anti-rabbit Alexa Fluor 488^®^ (goat polyclonal, Invitrogen # A32723, diluted 1:200). LacZ positive cells were detected with beta galactosidase staining (abcam, # ab102534), while acetylcholine receptors were detected by a-bungarotoxin (Alexa Fluor 488^®^, Invitrogen # B13422). Nuclei were stained with 300 nM DAPI (Thermo Fisher Scientific, # D1306).

Images were acquired using a Nikon ECLIPSE TE-2000 microscope equipped with UV source and CoolSnap Photometrics CCD camera.

### 2.8 Live/Dead assay

Cell viability in 3D wet-spun constructs was assessed 1 and 3 days after polymerization, by the use of Cell stain Double Staining Kit (Sigma-Aldrich), which allows the simultaneous fluorescence staining of viable and death cells. Briefly, after incubation of constructs with Calcein-AM (viable cells) and Propidium Iodide (dead cells) solutions for 30 minutes at 37°C, live and dead cells were displayed and photographed with Nikon ECLIPSE 2000-TE fluorescence microscope.

### 2.9 Image analysis

The alignment of the differentiated muscle fibers was assessed using OrientationJ plugin for ImageJ/FIJI [30]. The local orientation and isotropic properties, such as coherency, are evaluated for every pixel of the image based on structure tensors [31]. In order to analyze the images, we converted them in highly contrasted 8-bit images and processed them using OrientationJ dominant direction and OrientationJ distribution tools. OrientationJ dominant direction computes the coherency value, an index ranging between 0 and 1, with 1 indicating highly oriented structures and 0 indicating isotropic areas. OrientationJ distribution processes a weighted histogram of orientations, where the weight is the coherency itself.

### 2.10 Vessel density evaluation

To obtain a quantification of the vascularization levels in the normal muscle tissue and in the engrafted tissue, images from immunofluorescence analysis against SMA (Smooth Muscle Actine) and vW (von Willebrand factor), well-known vascularity markers, were analyzed using ImageJ program. The average fluorescence intensity of each immunofluorescence image was calculated as Mean Grey Value (*Analyze>Set Measurement>Mean Grey Value*, then *Analyze>Measure’*), an index elaborated by the program by assigning to each pixel a grey scale value. The sum of the gray values of all the pixels in the image are then divided by the total number of pixels. As a result, the higher the Mean Grey Value, the higher the average intensity of the fluorescent signal in the image. The total SMA-positive area was calculated on thresholded images (*Image>Adjust>Threshold*) using the command *Analyze Particles* and represents the total number of pixels that are part of the selection, spatially calibrated in squared μm *(Analyze>Set Scale)*. The results are indicative for the SMA expression extent in the tissue. The percentage of the SMA-positive area was calculated on thresholded images (*Image>Adjust>Threshold*) using the command *Analyze Particles* and represents the percentage of pixels that are part of the selection, indicative for the SMA expression extent in the tissue.

### 2.11 Statistical analysis

All experiments were performed in biological triplicate. Data were analyzed using GraphPad Prism 7, and values were expressed as means ± standard error (SEM). Statistical significance was tested using Student’s t-test. A probability of less than 5% (P < 0 05) was considered to be statistically significant.

## 3. Results

### 3.1 Assembly and characterization of a wet spinning set-up

Nowadays, biofabrication technologies hold great promises for the *in vitro* production of functional tissue equivalents thanks to their flexibility and unprecedent precision in cell and biomaterial deposition. Such features are of paramount importance for the recapitulation of organ/tissue complexity and functionality; especially true for the skeletal muscle, a tissue characterized by a hierarchical, uniaxially aligned architecture [32].

In a recent study, we presented an extrusion-based 3D bioprinting approach that enables to biofabricate hydrogel scaffolds composed of aligned myoblast-laden fibers [7]. Despite the remarkable results in terms of muscle architectural guidance, as shown by the accomplishment of oriented, mature myofibers that spontaneously contract *in vitro*, the developed approach suffers from several limitations. Firstly, ectopic implantation in a mouse model revealed some limitations in terms of scaffold remodeling, as large areas were actually devoid of myotubes (likely because of the large diameter of printed fibers ≈ 250 μm). Furthermore, in order to carefully pack adjacent fibers as close as possible, printing speed was relatively low (≈ 4 mm·s^-1^), thus limiting the possibility of biofabricating large constructs in an acceptable time.

In order to overcome these constraints, we have developed a custom-made wet-spinning system to generate well-organized myo-substitute with higher resolution (fiber diameter down to approx. 80 μm). Our novel wet-spinning system is notably simpler and easier to use then conventional 3D printers, as it comprises just a co-axial nozzle system and a Teflon drum connected to a stepper motor (**Figure 1a-c**). The highly aligned skeletal muscle architecture can be achieved by collecting the hydrogel fibers on the rotating drum, without any need of ad-hoc printing code. Additionally, the hydrogel fibers that are collected onto the drum form a highly packed bundle with almost no distance between adjacent fibers. This is hardly achievable with conventional extrusion bioprinting approaches (**Figure 1d,e**).

**Figure 1.**
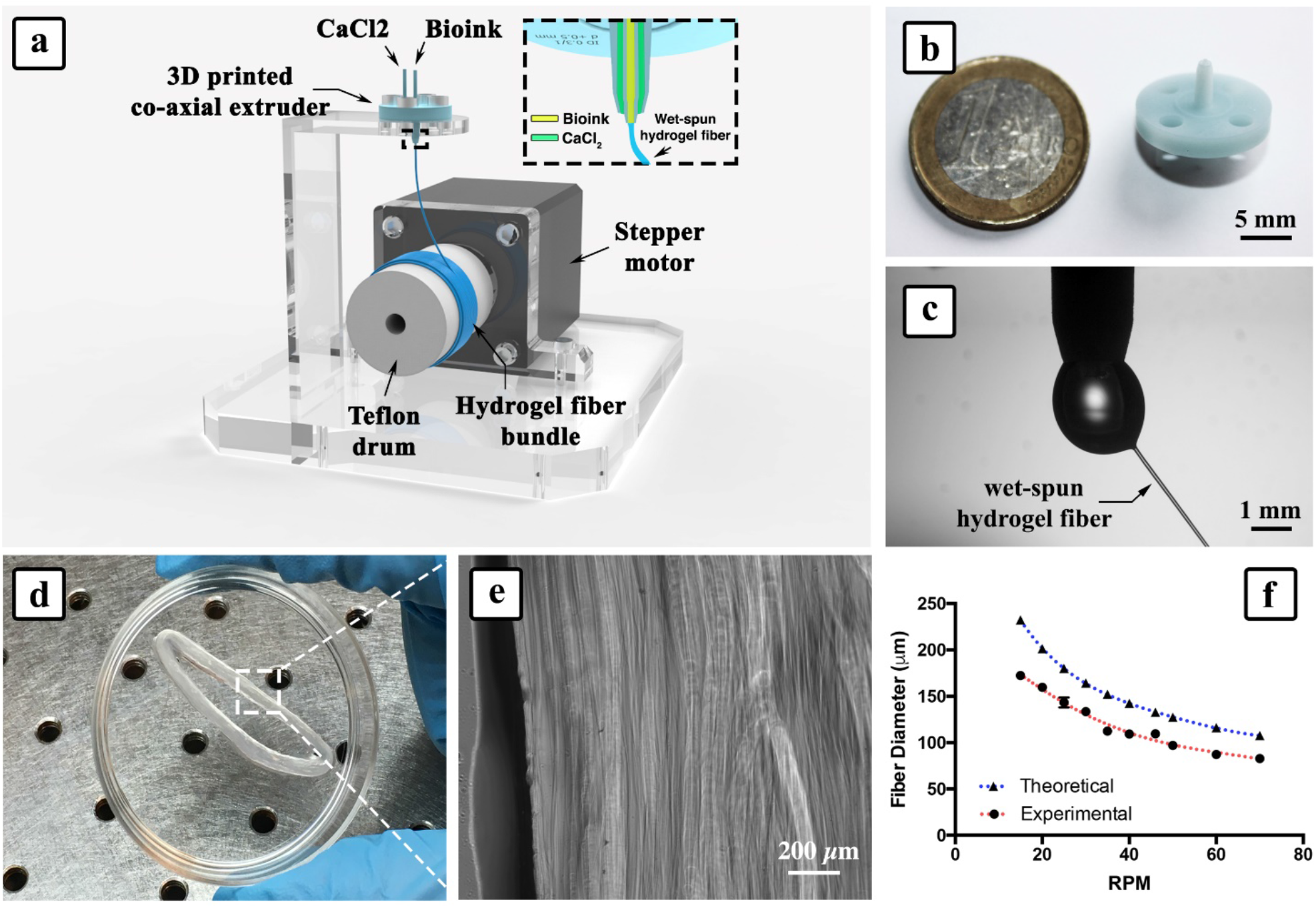
Wet-spinning set-up. a) Schematic representation of the wet-spinning set-up that was used for the biofabrication of myo-substitutes. b) 3D printed co-axial nozzle. c) Detail of the bioink extrusion at the tip of the co-axial needle. d) Hydrogel fiber bundle after removal from the drum and e) Bright field high magnification image showing densely packed fibers. f) Effect of motor speed on fiber diameter: interestingly, the experimental values are lower than the theoretical ones due to the shrinking of the alginate-based bioink upon exposure to divalent calcium ions.

Another advantageous feature of this system consists in the possibility to fine tune the hydrogel fiber size by controlling the flow rate of the bioink and the rotating speed of the drum. After a thorough characterization of the system, we found that hydrogel fibers can be produced in a reproducible manner using a motor speed in the 15÷70 rpm range (i.e. linear speed ≈ 20÷92 mm·s^-1^), which is 10-to 100-fold faster in comparison to conventional extrusion-based bioprinting approaches [33], and a bioink flow rate of 50 μL/min. As for the calcium chloride flow rate, we found that values in the range 15÷25 μL/min were equally suitable for fiber production, with no major impact on fiber size. As anticipated, the diameter of wet-spun fibers and the motor speed are inversely related (**Figure 1f**). The theoretical values of fiber diameters (*d_fiber_*) could be calculated by assuming a volumetric equivalence with the flow rate of the bioink (*Q_bioink_*) using the formula [27]:

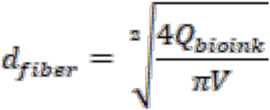

where *V* is the motor speed. Interestingly, such values were actually higher than the measured ones, for all motor speeds. Such effect was most likely a consequence of alginate shrinking upon exposure to calcium ions and was estimated to be of approximately 30%.

As bioink, we used a formulation already well established by our group containing PEG-Fibrinogen (PF), responsible for myogenic differentiation, and alginate (ALG), enabling immediate gelation of spun hydrogel fiber upon exposure to divalent calcium ions. This latter feature was exploited by using a co-axial nozzle extrusion system that allows to supply simultaneously the bioink and the primary crosslinking solution (CaCl_2_).

### 3.2 Biofabrication of highly-aligned muscle substitutes loaded with murine mesoangioblasts

After system optimization, we exploited the setup for the production of aligned yarns of cell-laden microfibers (**Figure 2**). In SMTE, next to matrix composition and scaffold architecture, the major role is played by the biological component, i.e. myogenic precursors. In this study, we initially fabricated hydrogel fiber yarns loaded with mouse-derived perivascular progenitor cells, mesoangioblasts (Mabs), and tested them for generation of skeletal muscle both *in vitro* and *in vivo*.

**Figure 2.**
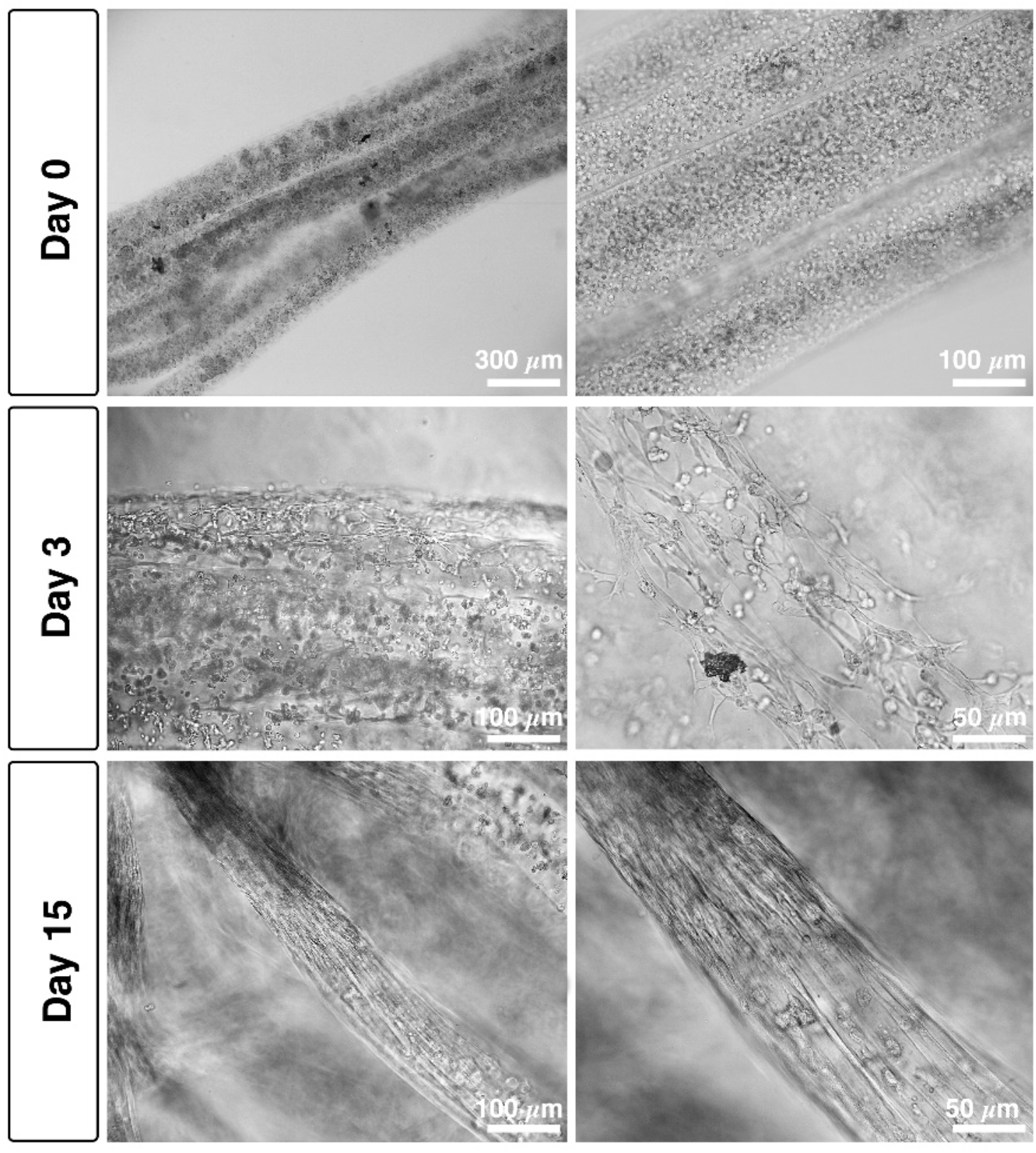
*In vitro* culture of Mabs-loaded hydrogel. Mabs-laden wet-spun yarns showing progressive myogenic differentiation over culturing time. As can be noted, Mabs morphology evolves from a round shape (single cells at early time points) into a more elongated one (generally within the first 3-5 days of culture), eventually forming long-range myotubes.

Mabs represent a primary myogenic cell population extensively characterized for its capability to robustly differentiate into skeletal muscle *in vitro* and *in vivo* [34]; human Mabs, have been employed in a clinical trial for Duchenne Muscular Dystrophy (Eudract Number: 2011-000176-33). As a preliminary test, we encapsulated Mabs within the fibers at different cell densities – ranging from 1 to 5×10^7^ cells/mL. As a result, we found that the best myogenic differentiation was obtained for concentration around 2÷2.5×10^7^ cells/mL (data not shown), a value in agreement with previously published studies [7]. As shown in **Figure 2**, Mabs had a typical round morphology upon encapsulation (day 0); cell elongation and spreading took place since the very first days of culture and was clearly appreciable at day 3 in vast areas of the engineered constructs. This was followed by an abundant myogenesis, with the formation of long-range, multi-nucleated myotubes within two weeks of culture (day 15). These myotubes were characterized by a remarkable aligned architectural organization.

To further validate our approach and especially its cell-compatibility towards primary cells, a live/dead staining based on calcein-AM (live cells in green) and propidium iodide (dead cells in red) was performed on the scaffolds at day 1 and 3. In both cases, the majority of Mabs appeared alive, with notable cell elongation at day 3 (see **Figure S1**), thus confirming the suitability of our approach for the biofabrication of myo-substitutes.

### 3.3 In vitro characterization of mouse-derived myo-substitutes

In order to assess the effectiveness and advantages of our wet-spinning system in the biofabrication of functional myo-substitute, we compared the *in vitro* myogenesis potential of Mabs in two hydrogel systems: a bulk gel and a wet-spun fiber yarn. The two systems are characterized by the same matrix composition and cell density but significantly different 3D architecture. *In vitro* myogenesis in the engineered myo-substitutes was evaluated by immunofluorescence staining against myosin heavy chain (MHC, a muscle-specific marker highly expressed in fully differentiated myotubes) after 20 days of culture (**Figure 3**). Mabs-laden yarns revealed a substantial myotube parallel organization and positivity for MHC (**Figure 3 upper panel**), while Mabs embedded in the bulk gels showed a similar MHC expression but an entangled, disordered myotube layout (**Figure 3 bottom panel**). Being a critical parameter for myo-substitutes functionality, myotube organization was further analyzed quantitatively through fluorescent image analysis. As shown in the polar plots in **Figure 3**, in Mabs-laden yarns all myotubes were distributed within ± 30° of average fiber orientation, whereas, in the case of bulk gels, myotube orientation was random with no preferential directions.

**Figure 3.**
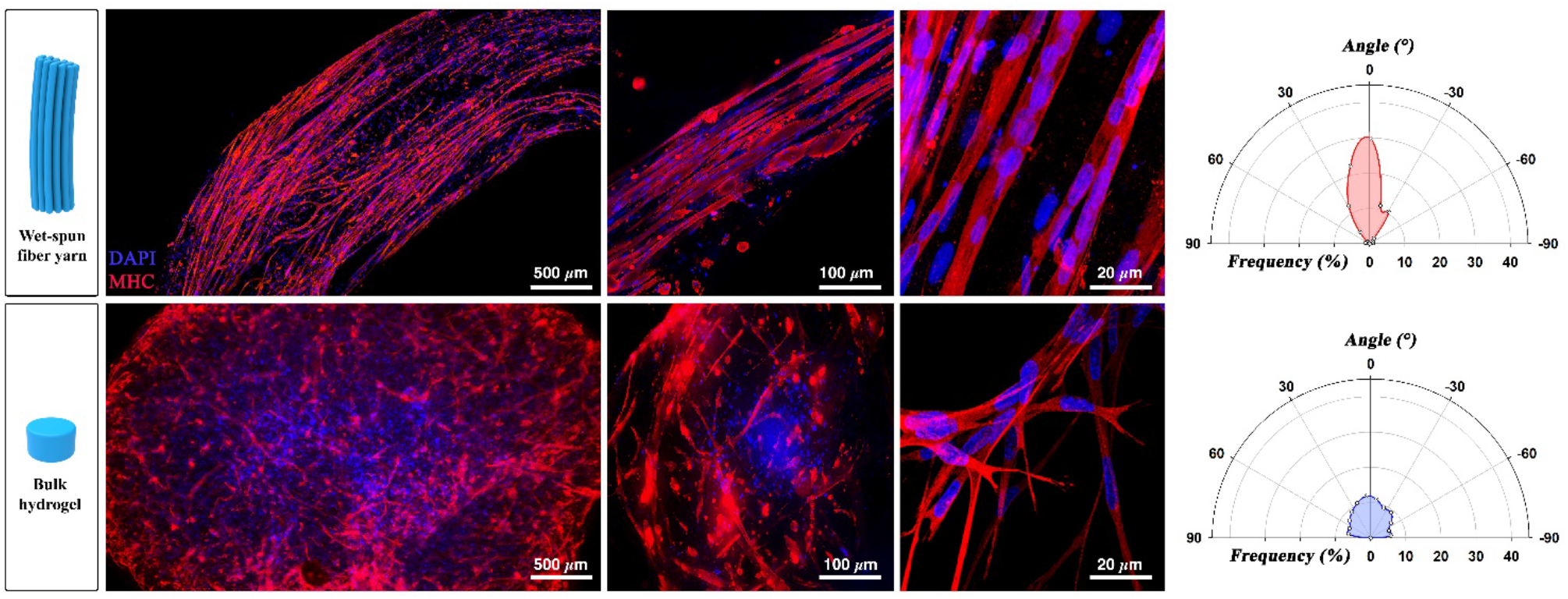
*In vitro* Mabs differentiation. MHC staining (red) performed on 3D printed and bulk structure loaded with Mabs after 15 days of culture reveal a significant difference in terms of myotube organization between the two hydrogel systems, with the wet-spun constructs greatly outperforming the bulk gels. Such difference – quantified by means of image analysis – is shown in the polar plots. Nuclei were counterstained by DAPI (Blue).

### 3.4 Tibialis Anterior (TA) regeneration by means of Mabs-derived myo-substitutes

Based on the promising *in vitro* results, Mabs-laden myo-substitutes were further tested *in vivo* in a volumetric muscle loss (VML) model following a surgical protocol previously established by our group [29]. The approach consists of surgically dislodging around 90% of the tibialis anterior muscle (TA) while leaving the surrounding tendons intact and in place (**Figure 4a**). In order to precisely distinguish Mabs-driven regeneration from that originating from host’s cells, the former were labelled by lentiviral transduction with nuclear β-Galactosidase (nLacZ) before grafting [29,34].

**Figure 4.**
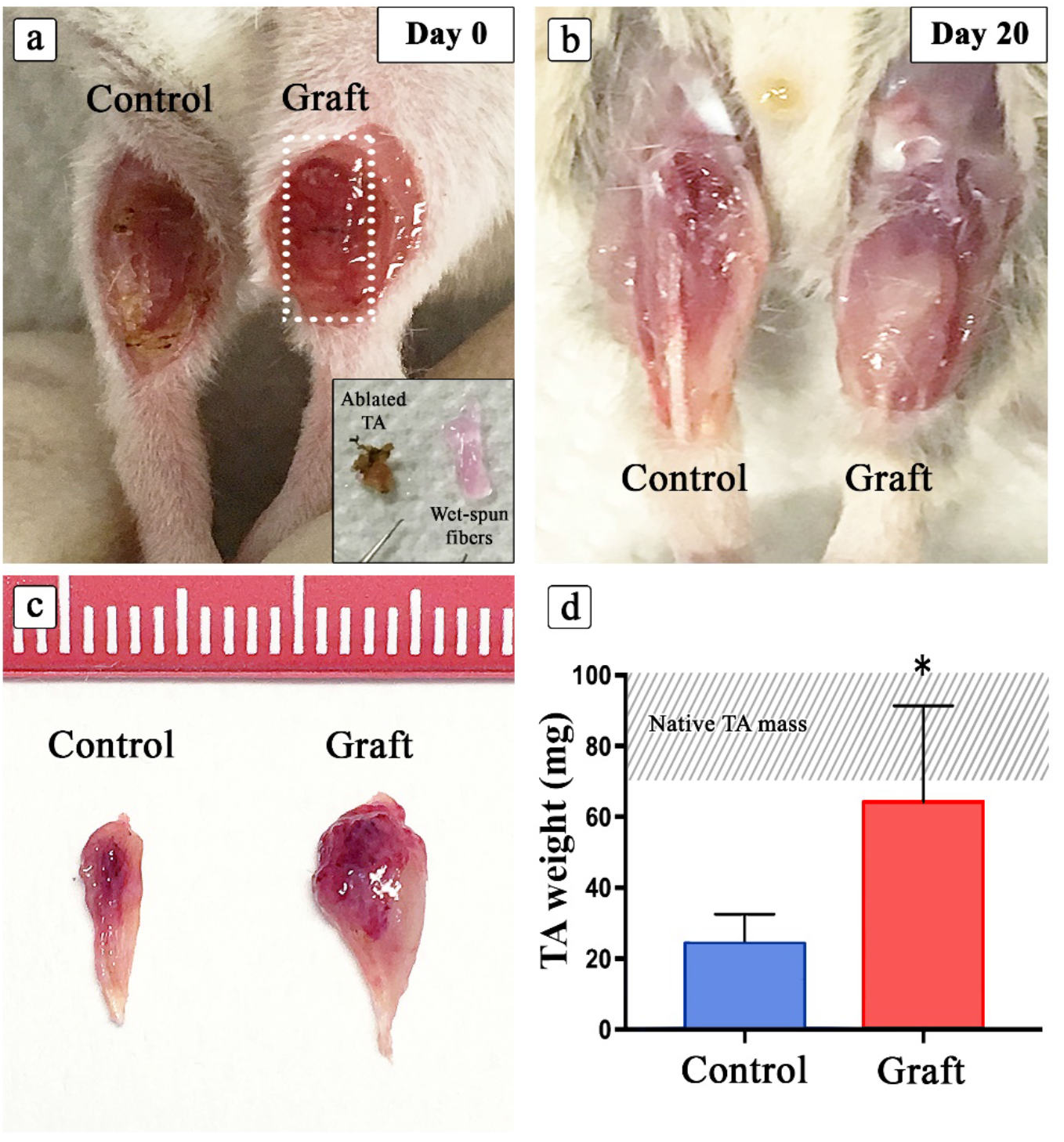
TA substitution by means of 3D wet-spun myo-substitute. a) Immunocompromised mouse leg showing ablated TA (control) and implanted TA (graft) at day 0; insert showing muscle mass removed next to Mabs loaded yarn. b) Implanted myo-substitute 20 days after grafting. c) Isolated TA from control ablated (control) and ablated/grafted leg (graft), the latter revealing a remarkable enhanced size. d) Graphical representation of the scored TA weight (n=6). The dashed area indicates the range of native TA mass depending primarily on mouse weight and age.

The results in the control animals (untreated) after 20 days post-implantation showed that, their leg presented a void where the TA was resected. Most of the missing muscular mass, was unable to self-regenerate after VML injury (**Figure 4b**). Conversely, wet-spun myo-substitute containing nLacZ-expressing Mabs implanted into the contralateral leg yielded a significant muscle regeneration whereby the TA defect was replenished by muscle-like tissue (**Figure 4b**). Furthermore, the recovered TA from the treated legs (implanted with wet-spun myo-substitutes) demonstrated larger mass compared to untreated control legs, as evidenced by the measured wet weights of both groups (**Figure 4c, d**).

Harvested tissues were further analyzed via histological staining to unveil their nature and anatomical structure. Transversal and longitudinal TA sections were firstly processed to detect nLacZ expression employing X-Gal substrate for β-Galactosidase activity. Next, samples were immunostained against MHC – for identification of muscle fibers – and laminin (LAM, basal lamina component surrounding muscle fibers), for architectural organization (**Figure 5**). As already revealed by macroscopic analysis, the cross-sectional area of the control and treated TAs present striking differences in terms of area. Importantly, this difference is predominantly associated with regeneration activity of the implanted Mabs as clearly evidenced by nLacZ staining (**Figure 5 upper panel**), showing a homogeneous distribution of Mabs throughout the crosssection of grafted leg. Moreover, the muscle architectural organization was almost completely reestablished as shown by the longitudinal cross-section displaying parallelly-oriented MHC positive myofibers surrounded by Laminin (**Figure 5 middle and bottom panels**). In addition to the reconstituted muscle structure, the magnified views of longitudinal sections underscore the typical hallmarks of sarcomerogenesis and then functional skeletal muscles (**Asterisks in Figure 5 bottom panels**). Accordingly, the disclosed sarcomeres were detected in close proximity to black X-Gal labeled nuclei, indicating the nLacZ/Mabs based myo-substitute were able to recapitulate all the morphological features of functional skeletal muscle tissue (**Arrows in Figure 5 bottom panels**). As widely demonstrated in our previous publication, acellular PF is completely inefficacious in VML TA mouse model, hence we prevent this further control avoiding unnecessary waste of animals [29].

**Figure 5.**
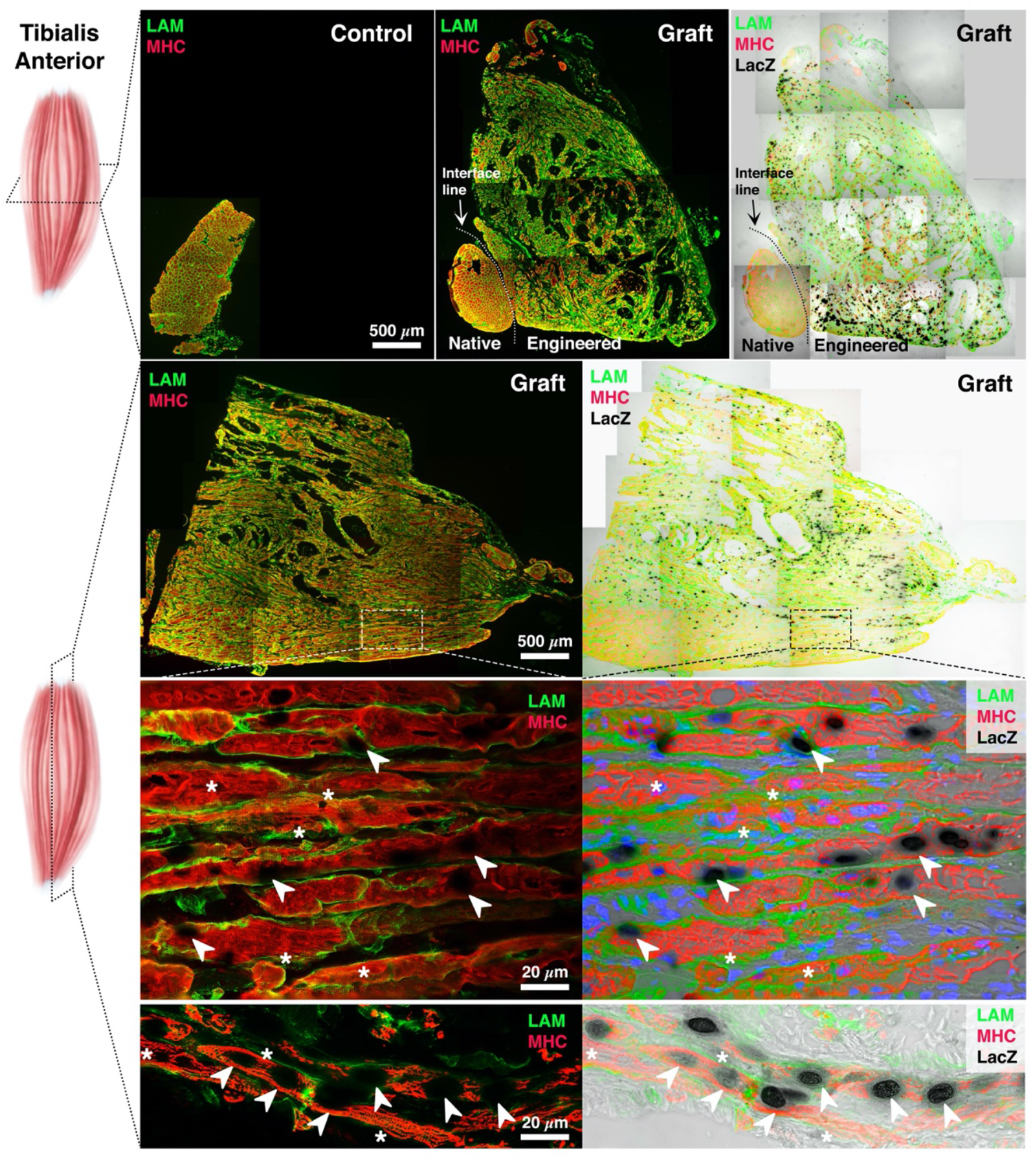
Immunofluorescence analysis on ablated TA muscle sections. Cross and longitudinal sections of ablated (control) and implanted (graft) TA were marked with LacZ and immunoassayed for MHC (red), LAM (green). Cross- and longitudinal sections were immunostained against MHC and LAM and overlaid to LacZ labeled images. Dashed area indicates the enlarged view displaying sarcomeres (asterisks) in combination with LacZ positive nuclei (arrowheads). Regenerated TA derived from nLacZ/Mabs-based myo-substitutes showed neo-vascularization – a key feature for artificial tissue subsistence and engraftment.

In addition to the muscle-like architectures there were also uncharacteristic gaps within the muscle tissue, these being noticeable as black voids in the cross-sections of the grafted TA. We speculate that these voids are due to the presence of some alginate residues from wet-spun fibers within the engineered muscle, which in physiological conditions degrades slowly and exclusively through the gradual exchange of calcium with monovalent ions [35]. Although this would appear as imperfections in the muscle repair, we hypothesize that such alginate could also exert a beneficial effect in the grafted muscle by acting as pillars, thus rapidly guiding myotube organization around them and providing additional mechanical support to the neo-tissue. Moreover, because of the small fiber size, such gaps do not significantly impair the functionality of the muscle as demonstrated by absolute force measurements of the muscle contraction (see **Figure 7**).

By immunolabelling the explanted grafts for von Willebrand factor (vW, specifically marking endothelial cells) and dystrophin (Dys, a key protein located in proximity of myotube membranes essential for proper muscle functionality, **Figure 6a**), it was possible to demonstrate that myotubes containing LacZ positive nuclei were surrounded by small vessel and capillaries (**Figure 6b-e**). In order to quantify vessel density in the reconstructed TA side, normal muscle tissue (native tissue) and reconstructed TA portion (graft) has been compared for vW fluorescent signal area and intensity by ImageJ analysis (**Supplementary Figure 2**). The evaluation revealed no statistically significant differences among the analyzed muscle moiety, further confirming the good vascularization of the reconstructed TA (**Figure 6f**). According to, vessel caliber and numbers were also evaluated by smooth muscle actin (SMA) vessel labelling marker. Despite an apparent larger area of the vessels in the normal muscle tissue (native) due to the greater caliber, in the reconstructed TA portion the number of the scored vessels is remarkably higher, confirming vW results and revealing a more than satisfactory vascularization of the reconstructed TA (**Supplementary Figure 3**).

**Figure 6.**
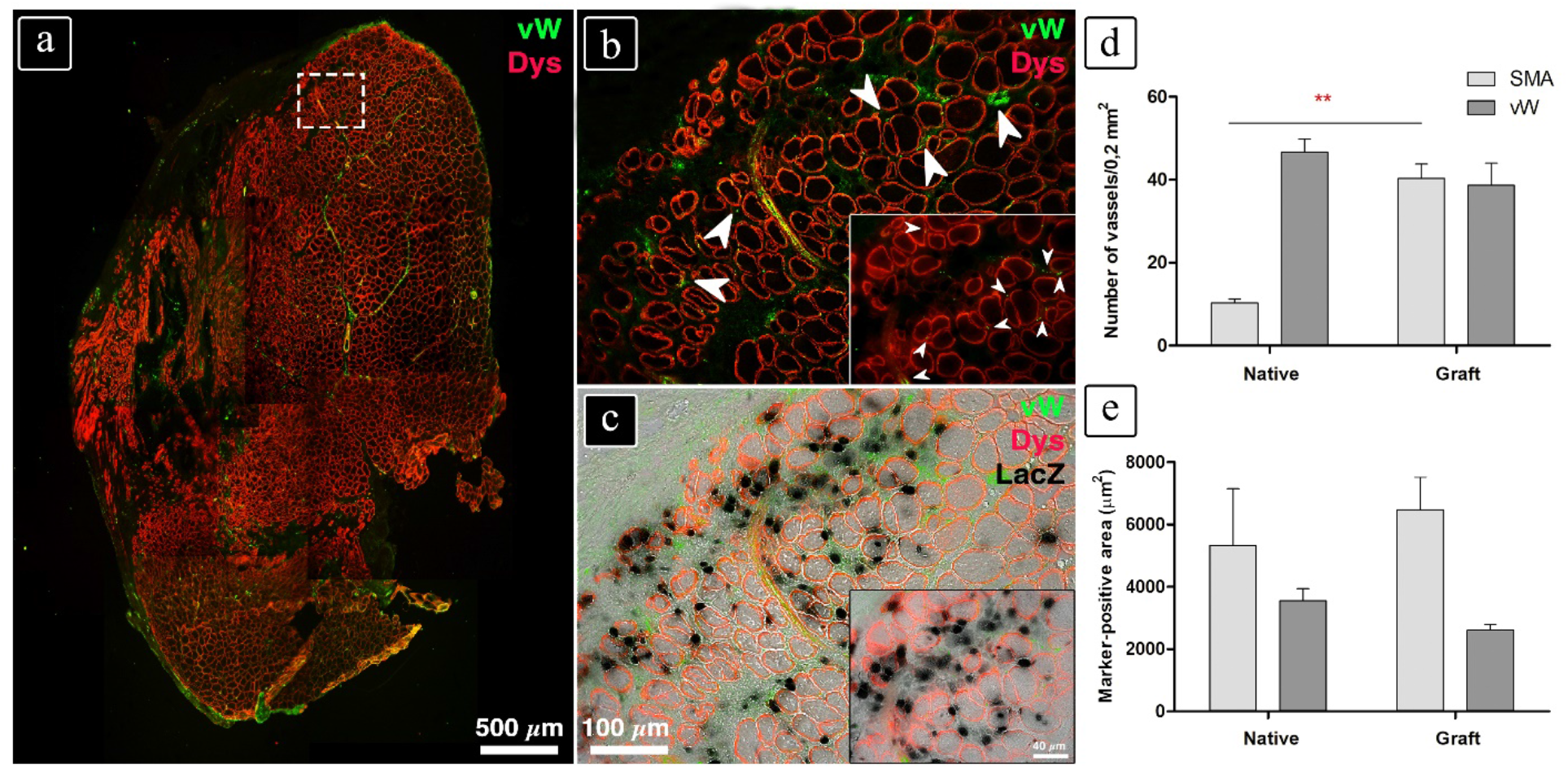
Neo-vascularization on TA sections implanted with nLacZ/Mabs derived myo-substitutes. a) TA reconstruction from immunofluorescence against vW (green) and Dys (red), white arrows indicating new vessel. b) Enlarged view of dashed area in (a). c) Superimposed image in (b) over LacZ stained image. d) Enlarged view of (b) image, capillaries pointed by white arrows. e) Superimposed image in (d) over LacZ stained image. f) Vessel density assessment by means of fluorescence quantitative evaluation, showing no significative differences between normal muscle (native tissue) and implanted one (graft), (n=3).

In addition to the morphological analysis, we also carried out a functional assessment of the harvested engineered muscle to evaluate TA muscular activity following massive ablation. Muscle innervation is a necessary condition for ensuring voluntary contraction activation and functionality. Hence, we investigated nerve supply by means of immuno-identification of axons and neuromuscular junctions into engineered muscle fibers (**Figure 7**). Staining against anti phospho-neurofilament (NeuF) has been employed to identify motor neuron axons approaching the myofibers. As shown in **Figure 7a** (marked with an asterisk), NeuF positive axons inserted between MHC positive myofibers innervating the engineered TA derived from nLaz/Mabs (**Figure 7b**). Additionally, simultaneous staining against a-Bungarotoxin (BTX – used to label the acetylcholine receptor in the chemical synapses) and NeuF was used to unveil the formation of numerous neuromuscular junctions (NMJ), with BTX and NeuF pointing out the postsynaptic membrane laying on the muscle fibers and the axon presynaptic cell residing in between myofibers respectively (**arrows in Figure 7c**). In particular, detailed view of the NMJ from reconstructed TA in the grafted leg reveals the characteristic pretzel-like structure positive for BTX with axonal inserts being positive for NeuF (**inset in Figure 7c**).

**Figure 7.**
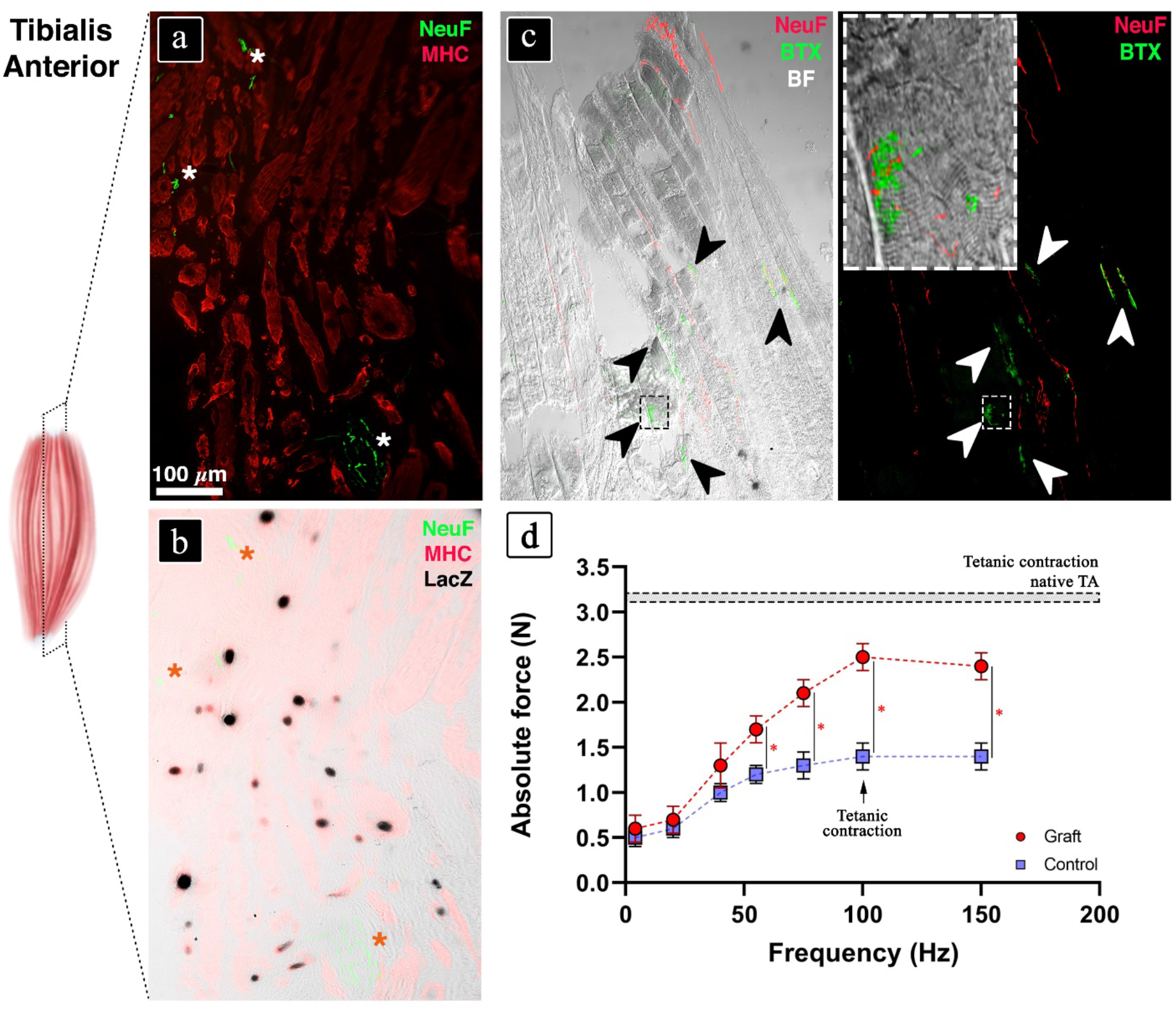
Innervation assessment on reconstructed TA sections. a) Immunofluorescence against NeuF (green) and MHC (red) revealing reconstructed muscle innervation (asterisks). b) Superimposed image in (a) over LacZ stained image. c) NeuF and BTX immunolabelled sections showing neuromuscular junction formation (arrowheads), insert enlarged view of dashed box. d) Electrophysiological analysis scoring muscle strength recovery performed *in vivo* on ablated TA (control) and wet-spun implanted contralateral TA (graft) (n=3).

To further confirm the functionality of the reconstructed TA, we performed *in vivo* electrophysiological analysis using a well-established experimental set up used for measuring the actual force production (see **Supplementary Figure 2** and **Supplementary Table 1**). [36]. The testing consisted of a direct electrical stimulation of the common peroneal nerve and the simultaneous measurement of the force generated by the TA. The results demonstrated a higher force production in the engineered TA compared to the control, reaching almost a two-fold increase of tetanic force for frequencies around 100 Hz (**Figure 7d**).

Besides all the above analyzed parameters demonstrating the completeness of the reconstructed TA in terms of histoarchitecture and functionality, we have not leaved out the replenishment of stem cell niche. Hence, we investigated by means of immunofluorescence the presence of Pax-7 positive muscle resident stem cells, namely satellite cells, in the engineered TA muscle. Notably, we identified satellite cells in between LacZ positive myotubes (arrowheads in **Supplementary Figure 3**), revealing also the provision of myogenic stem cells in the reconstructed mouse TA. Taken together, the morphological and functional characterization of the engineered TA reveal an unprecedented reconstructive capability of our system, resulting in a swift recovery of the VML defect. The fact that the engineered TA recovered partial functionality only 20 days after implantation further underscores the remarkable speed of the muscle recovery after such a traumatic injury.

### 3.5 In vitro characterization of human-derived myo-substitutes

Mouse derived myo-structures showed a remarkable capacity in generating functional myo-substitutes *in vitro* and *in vivo*. However, in order to eventually translate such technology to a clinical scenario, the performance of the proposed biofabrication approach in combination with human myogenic progenitors needs to be assessed. Stem cells stability and functionality represent a key task for fabrication of large 3D structures [37]. Mechanical forces acting on cells during bioink extrusion – such as high shear-stresses – have been demonstrated to directly impact cell proliferation and cell lineage commitment [38,39].

Moreover, different cell lineages, namely mouse Mabs and human primary myoblast, could react differently to the bioink mechano-chemical properties as extensively demonstrated for other myogenic stem/progenitor cells [40,41]. Hence, we performed a preliminary set of *in vitro* experiments with the sole intent of demonstrating the compatibility of the proposed approach with primary human myoblasts (hMyob) and verifying cell stability in terms of myogenic differentiation capacity. To this aim, we benchmarked – as in the case of Mabs – the myo-substitutes fabricated through the proposed wet-spinning technology with standard bulk gel preparations (**Figure 8**).

**Figure 8.**
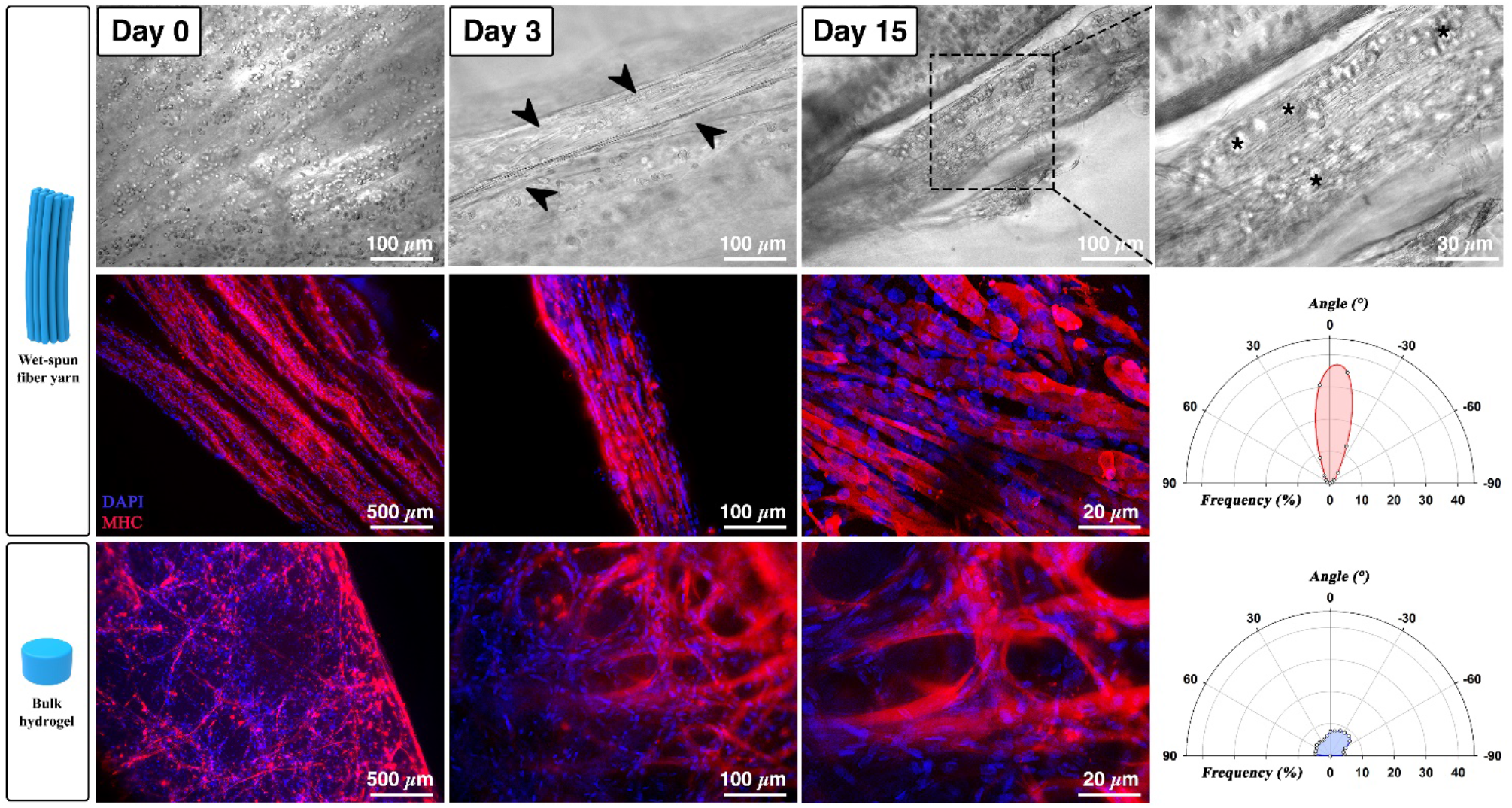
Fabrication of human myoblast based myo-substitute. Wet-spun yarn containing human myogenic precursors showed progressive cell differentiation with the time (arrows in day 3); dashed box enlargement reveals still undifferentiated cells after 15 days of culture (Upper panel). Middle and lower panel showing MHC immunostaining (red) performed on microfiber yarns and bulk structure loaded with human myoblasts upon 15 days of culture, revealing a significant difference in terms of myotube organization between the two hydrogel systems, with the wet-spun constructs greatly outperforming the bulk gels. Such difference is quantified in the polar plots. Nuclei were counterstained by DAPI (Blue).

The hMyob confined into the tiny fibers underwent myogenic differentiation following a timeline similar to murine Mabs, with a pronounced cell elongation appreciable as early as day 3 (see black arrows in **Figure 8**, day 3) and an abundant formation of myotube bundles around day 15 of culture (see magnified image of day 15 in **Figure 8**). The morphological analysis performed 15 days after fabrication by MF20 immunolabelling revealed a substantial myogenic differentiation in both hydrogel systems with an abundant expression of MHC. Also in this case, a significant improvement in the 3D organization of the forming myotubes is noticed in the comparison between the wet-spun fiber yarn and the bulk gel system. As confirmed by the quantitative analysis of myotube orientation shown in the polar plots in **Figure 8** the former clearly outperforms the latter.

## 4. Discussion

The *in vitro* fabrication of human tissue equivalents and the *in vivo* restoration of tissue/organ functionalities is one of the biggest biomedical challenges of our times. Thanks to the biotechnological progress of the last two decades, substantial advances in the tissue engineering field have been achieved with the fabrication of liver tissue with sinusoids [42,43], lung functions [44], spleen [45], skeletal muscle model functionality [46,47] *in vitro* models that are capable of mimicking native organ morphology and functionalities.

In the specific context of skeletal muscle tissue engineering (SMTE), researchers have greatly improved myo-substitute 3D morphological organization and functionality by i) developing specific dynamic culturing protocols, which generally include an electro-mechanical stimulation of the engineered samples, ii) implementing advanced technologies such as 3D bioprinting or organs-on-a-chip, and iii) supplying bioactive molecules in combination with tailored matrices to promote myogenic differentiation.

A common limitation shared by all the developed systems consists in the recapitulation of skeletal muscle histoarchitecture and functionality to a limited scale – generally from hundreds of micrometers to a few millimeters. Despite being sufficient for all those applications in which the overall size of the engineered construct does not influence the quality and reliability of the obtained results – such as in the case of miniaturized biohybrid actuators or simplified skeletal muscle models, such construct volumes are far from those needed in a clinical scenario (up to hundreds of cm^3^). Therefore, the availability of a system enabling the fabrication of macroscopic myo-substitutes that could eventually be translated into the clinics is still an unmet need.

Over the past decade, a condition clinically referred to as volumetric muscle loss (VML), has been the subject of a number of studies. A variety of approaches including acellular gels [48], cell sheet-derived [49], minced muscle [50] or tissue engineering grafts [29] have been proposed for VML treatment. Among these strategies, as described in a recent excellent review on this topic [20], the use of grafts delivered in combination with a myogenic cell source should be preferred to the other methods. In fact, acellular gels, as ultimately demonstrated in a recent study by Corona [51] do not offer any beneficial effect for VML recovery. Other strategies suffer from major drawbacks as well, such as scalability, difficulty of handling/surgical implantation (especially in the case of cell sheet) and limited volume of muscle tissue harvestable from the patient. All of these drawbacks complicate the clinical translation for such approaches. On the other hand, tissue engineered muscle grafts (TEMGs) offer the advantages of enhanced construct durability, ease of surgical manipulation and high scalability potential. However, due to the lack of a standard reference VML model, comparison between studies is not trivial due to the broad range of myogenic precursors and scaffold materials as well as differences in *in vitro* pre-culturing time, construct size, and type of VML model – i.e. animal used, size of VML defect, muscle anatomy and pennation. Therefore, in order to compare more effectively our findings with the state-of-the-art, we have selected only those studies in which a VML defect was created in the tibialis anterior of small size animals – i.e. mice and rats. A brief overview of the results described in these studies is reported in **Table 1**.

**Table 1.**
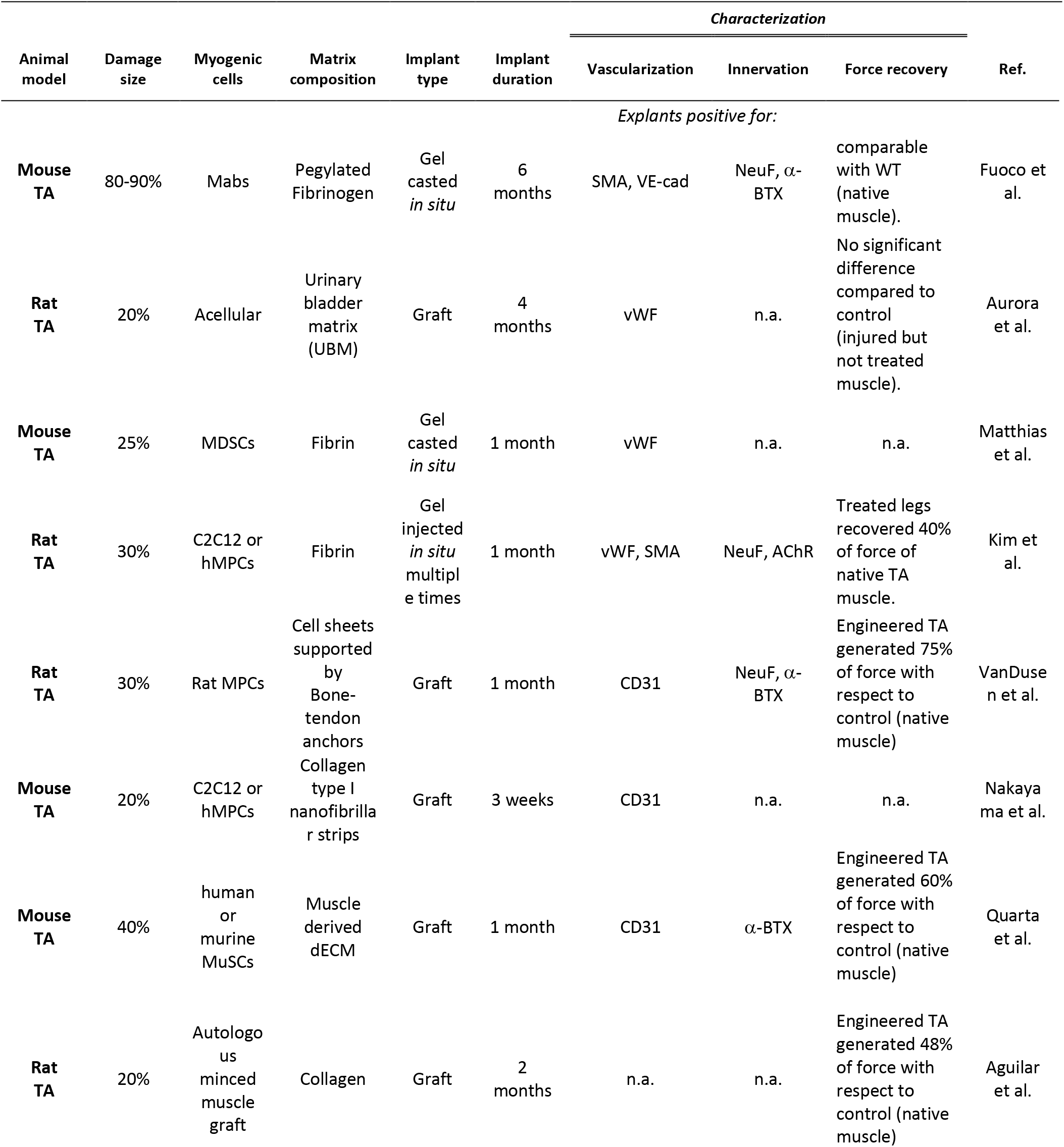
State-of-the-art of TA-VML reconstruction in small size animals (mice and rats).

As can be observed in the table, a different VML model was employed in each study with the damage size ranging generally between 20-40% of TA volume (with the exception of the study from Fuoco et al. in which the VML defect size reached approx. 90% of the TA volume); implant duration then range from 3 weeks to 6 months. Such variability of VML models – compounded by the use of different scaffold materials and myogenic precursors – makes it difficult to rank the different approaches. Nevertheless, there are a few outcomes that stand out. For example, the worst results in terms of distribution of force recovery were obtained for acellular implants (no significant difference compared to control leg). The most promising results were achieved with a PEGylated Fibrinogen gel loaded with Mabs casted directly *in situ* (force comparable with WT native muscle) 6 months after implantation.

Despite the promising outcomes in force and muscle mass recovery, the obtained results are still unsatisfactory due to the recurring problems related to cell engraftment, differentiation efficacy, muscle fibrosis, long regeneration time (up to 6 months) and limited system scalability [8,9,48]. Scalability – being a pivotal requirement for translating animal model approaches into human clinical therapy – is a critical feature that has so far often limited the possibility to translate the aforementioned approaches into VML models of large size animals [51].

In this context, our work proposes an easily scalable SMTE approach, which is completely independent from muscle defect geometry, based on a simple yet robust biofabrication wet-spinning strategy that enables the assembly of large myo-substitutes capable of efficiently guiding and speeding up cell differentiation and 3D muscle tissue organization both *in vitro* and *in vivo*. These features are successfully achieved by assembling highly anisotropic bundles composed of tiny hydrogel fibers where myogenic precursors are geometrically confined. In spite of their simplicity, such architectures cannot be fabricated using other fiber-based technique – e.g. 3D bioprinting – due to technical limitations such as printing speed and resolution [7].

In our previous work, we have reported an advanced reconstruction of a functional mouse TA [29]; nevertheless, a long implantation time (6 months) was needed to reestablish TA mass, morphology and functionality. Moreover, the proposed strategy relied on the polymerization *in situ* of a liquid hydrogel-based precursor solution – a strategy suitable for the TA lodge thanks to its concave shape but hardly transferable to other muscle locations.

Here, we have demonstrated that the aforementioned limits can be overcome by employing a biofabricated wet-spun fiber yarn loaded with primary muscle precursors. Such constructs, in fact, can be easily manipulated, thus being totally independent from the transplantation muscle shape and size. Moreover, these constructs have shown an extraordinary capacity in regenerating muscle mass, vasculature and innervation thus allowing recovery of muscle functions within only 20 days after implantation upon a massive TA ablation (~ 90% muscle mass removed). Last, but definitely not least, the preliminary evidences of compatibility of our approach with human primary myoblast open a practical possibility to translate this approach into a clinical scenario.

Currently, we are investing a great deal of work to improve construct matrix remodeling *in vivo* by precisely tuning bioink composition and to incorporate vasculature and innervation features during the biofabrication process. The latter aspects, in fact, are undoubtedly needed to successfully translate our approach into large animal VML models. In addition, we are planning to test our approach for VML recovery in a swine model using autologous cells.

In conclusion, in light of the presented results, we think that the proposed strategy represents a breakthrough in the field and, with further refinements, may be successfully translated to the clinic as a reconstructive therapy for VML patients.

### The paper explained

#### Problem

Skeletal muscle regenerative capability is related to damage size, in other words small injuries are able to self-repair while larger lesions need therapeutic treatment and/or surgical intervention. Muscular dystrophies and large trauma due to both accidental tearing or cancer ablation surgery lead to extensive loss of tissue, requiring a reconstructive approach to recover the muscle damage. Within this scenario skeletal muscle tissue engineering represent a real opportunity to replace damaged or lost muscular tissue with complete and functional implantable artificial muscle. But in spite of everything there is still an open challenge related to the architectural muscular organization replica and overall the functionality in terms of vessel and nerves supply.

#### Results

Exploiting homemade 3D printing technology to assemble in orderly way myogenic precursor cells and supporting matrix, we were able to replace large damaged muscle mass and rapidly restore (within 3 weeks) muscular histoarchitecture and functionality in a mouse model of extensive muscle damage. Forcing parallel alignment by 3D printing innovative approach, macroscopic artificial muscular structures were generated *in vitro*. Upon implantation, the printed muscle substitutes supported the formation of new blood vessels and neuromuscular junctions, fundamental for artificial muscle survival and functional recovery.

#### Impact

These data represent a solid base for further testing the 3D printing technology here proposed in large animal size, before to eventually be translated into clinical scenarios for the treatment of a wide range of pathology eliciting muscle wasting.

## Acknowledgments

We would like to thanks Giulio Cossu for critical reading. This study was supported by National Science Centre Poland (NCN) within SONATA 14 project No. 2018/31/D/ST8/03647 to MC, Italian Ministry of University and Research PRIN funding scheme No. 2015FBNB5Y_002 to CG and 201742SBXA_004 to SC.

## Author contributions

MC and CG conceived and designed the experiments; MC, ST, EF, and CF carried out experiments; CG, SC and GC supervised the project. AR acquired fluorescence images relative to in vitro culture; MC, MN, WS, PG and ST conceived, designed and optimized the co-axial wet-spinning system; LV isolated and cultured human primary myoblast beside critical reading; JB, CZ and RB provided surgical support for TA muscle ablation; BB performed electrophysiological analysis together with critical reading; DS provided PEG-fibrinogen characterization and production next to critical reading. MC, ST, EF and CG wrote the manuscript. All co-authors contributed to discussion and analysis of the data.

## Supplementary Infromation

Fig. S1. Live/dead staining of yarns loaded with mouse primary cells (Mabs).

Fig. S2. Fluorescence image and relative ImageJ output analysis employed for vessel density evaluation.

Fig. S3. Vascularization assessment on TA reconstructed upon 3D wet-spun myo-substitute implantation.

Fig. S4. Electrophysiological set-up.

Fig. S5. Satellite cells Pax-7 labelling on reconstructed TA sections.

Table S1. Absolute force raw data obtained from electrophysiological set-up.

## References

1. Carlson BM (1973) The regeneration of skeletal muscle — a review. Am J Anat 137: 119–149.

2. Huard J, Li Y, Fu FH (2002) Muscle injuries and repair: current trends in research. J bone Jt surgery Am Vol 84: 822–832.

3. Chargé SBP, Rudnicki MA (2004) Cellular and Molecular Regulation of Muscle Regeneration. Physiol Rev 84: 209–238.

4. Rizzi R, Bearzi C, Mauretti A, Bernardini S, Cannata S, Gargioli C (2012) Tissue engineering for skeletal muscle regeneration. Muscles Ligaments Tendons J 2: 230–234.

5. Grogan B, Hsu J (2011) Volumetric muscle loss. Am Acad Orthop Surg 19: S35–37.

6. Mercuri E, Muntoni F (2013) Muscular dystrophies. Lancet 381: 845–860.

7. Costantini M, Testa S, Mozetic P, Barbetta A, Fuoco C, Fornetti E, Tamiro F, Bernardini S, Jaroszewicz J, Trombetta M, et al. (2017) Microfluidic-enhanced 3D bioprinting of aligned myoblast-laden hydrogels leads to functionally organized myofibers in vitro and in vivo. Biomaterials.

8. Fuoco C, Cannata S, Gargioli C (2016) Could a functional artificial skeletal muscle be useful in muscle wasting? Curr Opin Clin Nutr Metab Care 19: 182–187.

9. Fuoco C, Petrilli LL, Cannata S, Gargioli C (2016) Matrix scaffolding for stem cell guidance toward skeletal muscle tissue engineering. J Orthop Surg Res 1–8.

10. Caron L, Kher D, Leong Lee K, Mckernan R, Biljana D, Hidalgo A, Li J, Yang H, Main H, Ferri G, et al. (2016) A Human Pluripotent Stem Cell Model of Facioscapulohumeral Muscular Dystrophy-Affected Skeletal Muscles. Stem Cells Transl Med 5: 1145–1161.

11. Chal J, Tanoury Z Al, Hestin M, Gobert B, Aivio S, Hick A, Cherrier T, Nesmith AP, Parker KK, Pourquié O (2016) Generation of human muscle fibers and satellite-like cells from human pluripotent stem cells in vitro. Nat Protoc 11:.

12. Tanaka A, Woltjen K, Miyake K, Hotta A, Ikeya M, Yamamoto T (2013) Efficient and Reproducible Myogenic Differentiation from Human iPS Cells: Prospects for Modeling Miyoshi Myopathy In Vitro. PLoS One 8:.

13. Smith AST, Davis J, Lee G, Mack DL, Kim D (2016) Muscular dystrophy in a dish: engineered human skeletal muscle mimetics for disease modeling and drug discovery. Drug Discov Today 00: 1–12.

14. Ostrovidov S, Salehi S, Costantini M, Suthiwanich K, Ebrahimi M, Sadeghian RB, Fujie T, Shi X, Cannata S, Gargioli C, et al. (2019) 3D Bioprinting in Skeletal Muscle Tissue Engineering. Adv Sci News 1805530: 1–14.

15. Juhas M, Ye J, Bursac N (2016) Design, evaluation, and application of engineered skeletal muscle. Methods 99: 81–90.

16. Truskey GA, Achneck HE, Bursac N, Chan HF, Cheng CS, Fernandez C, Hong S, Jung Y, Koves T, Kraus WE, et al. (2013) Design considerations for an integrated microphysiological muscle tissue for drug and tissue toxicity testing. Stem Cell Res Ther 4: 1–5.

17. Agrawal G, Aung A, Varghese S (2017) Skeletal muscle-on-a-chip: An in vitro model to evaluate tissue formation and injury. Lab Chip 17: 3447–3461.

18. Leng L, McAllister A, Zhang B, Ranu A, Radisic M, Guenther A (2012) Osaic hydrogels: One-step formation of multiscale soft materials. Adv Mater 24: 3650–3658.

19. Zhang B, Montgomery M, Davenport-H L, Korolj A, Radisic M (2015) Platform technology for scalable assembly of instantaneously functional mosaic tissues. Sci Adv 1:.

20. Gilbert-Honick J, Grayson W (2019) Vascularized and Innervated Skeletal Muscle Tissue Engineering. Adv Healthc Mater 1900626: 1–27.

21. Passipieri JA, Christ GJ (2016) The potential of combination therapeutics for more complete repair of volumetric muscle loss injuries: The role of exogenous growth factors and/or progenitor cells in implantable skeletal muscle tissue engineering technologies. Cells Tissues Organs 202: 202–213.

22. Khouri RK, Cooley BC, Kunselman AR, Landis JR, Yeramian P, Ingram D, Natarajan N, Benes CO, Wallemark C, Anthony J, et al. (1998) A prospective study of microvascular free-flap surgery and outcome. Plast Reconstr Surg 102: 711–721.

23. Lin C-H, Wei F-C, Levin S, Chen M (1999) Donor-Site Morbidity Comparison between Endoscopically Assisted and Traditional Harvest of Free Latissimus Dorsi Muscle Flap. Plast Reconstr Surg 104: 1070–1077.

24. Machingal MA, Corona BT, Walters TJ, Kesireddy V, Koval CN, Dannahower A, Zhao W, Yoo JJ, Christ GJ (2011) A tissue-engineered muscle repair construct for functional restoration of an irrecoverable muscle injury in a murine model. Tissue Eng Part A 17: 2291–2303.

25. Rinoldi C, Costantini M, Kijeńska-Gawrońska E, Testa S, Fornetti E, Heljak M, Ćwiklińska M, Buda R, Baldi J, Cannata S, et al. (2019) Tendon Tissue Engineering: Effects of Mechanical and Biochemical Stimulation on Stem Cell Alignment on Cell-Laden Hydrogel Yarns. Adv Healthc Mater 8: 1–10.

26. Almany L, Seliktar D (2005) Biosynthetic hydrogel scaffolds made from fibrinogen and polyethylene glycol for 3D cell cultures. Biomaterials 26: 2467–2477.

27. Colosi C, Costantini M, Latini R, Ciccarelli S, Stampella A, Barbetta A, Massimi M, Conti Devirgiliis L, Dentini M (2014) Rapid prototyping of chitosan-coated alginate scaffolds through the use of a 3D fiber deposition technique. J Mater Chem B 2: 6779–6791.

28. Vianello S, Pantic B, Fusto A, Bello L, Galletta E, Borgia D, Gavassini BF, Semplicini C, Sorarù G, Vitiello L, et al. (2017) SPP1 genotype and glucocorticoid treatment modify osteopontin expression in Duchenne muscular dystrophy cells. Hum Mol Genet 26: 3342–3351.

29. Fuoco C, Rizzi R, Biondo A, Longa E, Mascaro A, Shapira-Schweitzer K, Kossovar O, Benedetti S, Salvatori ML, Santoleri S, et al. (2015) In vivo generation of a mature and functional artificial skeletal muscle. EMBO Mol Med 7: 411–422.

30. Püspöki Z, Storath M, Sage D, Unser M (2016) Transforms and Operators for Directional Bioimage Analysis: A Survey.

31. Rezakhaniha R, Agianniotis A, Schrauwen JTC, Griffa A, Sage D, Bouten CVC, Van De Vosse FN, Unser M, Stergiopulos N (2012) Experimental investigation of collagen waviness and orientation in the arterial adventitia using confocal laser scanning microscopy. Biomech Model Mechanobiol 11: 461–473.

32. Frontera WR, Ochala J (2015) Skeletal Muscle: A Brief Review of Structure and Function. Calcif Tissue Int 45: 183–195.

33. Jian H, Wang M, Wang S, Wang A, Bai S (2018) 3D bioprinting for cell culture and tissue fabrication. Bio-Design Manuf 1: 45–61.

34. Fuoco C, Salvatori M, Biondo A, Shapira-Schweitzer K, Santoleri S, Antonini S, Bernardini S, Tedesco FS, Cannata S, Seliktar D, et al. (2012) Injectable polyethylene glycol-fibrinogen hydrogel adjuvant improves survival and differentiation of transplanted mesoangioblasts in acute and chronic skeletal-muscle degeneration. Skelet Muscle 2: 24.

35. Kong HJ, Kaigler D, Kim K, Mooney DJ (2004) Controlling rigidity and degradation of alginate hydrogels via molecular weight distribution. Biomacromolecules 5: 1720–1727.

36. Blaauw B, Canato M, Agatea L, Toniolo L, Mammucari C, Masiero E, Abraham R, Sandri M, Schiaffino S, Reggiani C (2009) Inducible activation of Akt increases skeletal muscle mass and force without satellite cell activation. FASEB J 23: 3896–3905.

37. Kang H-W, Lee SJ, Ko IK, Kengla C, Yoo JJ, Atala A (2016) A 3D bioprinting system to produce human-scale tissue constructs with structural integrity. Nat Biotechnol 34: 312–319.

38. Engler AJ, Sen S, Sweeney HL, Discher DE (2006) Matrix elasticity directs stem cell lineage specification. Cell 126: 677–689.

39. Bianco P, Robey PG (2001) Stem cells in tissue engineering. Nature 414: 118–121.

40. Gilbert PM, Havenstrite KL, Magnusson KEG, Sacco A, Leonardi NA, Kraft P, Nguyen NK, Thrun S, Lutolf MP, Blau HM (2010) Substrate elasticity regulates skeletal muscle stem cell self-renewal in culture. Science (80-) 329: 1078–1081.

41. Nakayama KH, Shayan M, Huang NF (2019) Engineering Biomimetic Materials for Skeletal Muscle Repair and Regeneration. Adv Healthc Mater 8: 1–14.

42. Schütte J, Hagmeyer B, Holzner F, Kubon M, Werner S, Freudigmann C, Benz K, Böttger J, Gebhardt R, Becker H, et al. (2011) ‘Artificial micro organs’ -A microfluidic device for dielectrophoretic assembly of liver sinusoids. Biomed Microdevices 13: 493–501.

43. Toh YC, Lim TC, Tai D, Xiao G, Van Noort D, Yu H (2009) A microfluidic 3D hepatocyte chip for drug toxicity testing. Lab Chip 9: 2026–2035.

44. Huh D, Matthews BD, Mammoto A, Montoya-Zavala M, Yuan Hsin H, Ingber DE (2010) Reconstituting organ-level lung functions on a chip. Science (80-) 328: 1662–1668.

45. Baker M (2011) Tissue models: A living system on a chip. Nature 471: 661–665.

46. Nagamine K, Kawashima T, Sekine S, Ido Y, Kanzaki M, Nishizawa M (2011) Spatiotemporally controlled contraction of micropatterned skeletal muscle cells on a hydrogel sheet. Lab Chip 11: 513–517.

47. Serena E, Zatti S, Zoso A, Lo Verso F, Tedesco FS, Cossu G, Elvassore N (2016) Skeletal Muscle Differentiation on a Chip Shows Human Donor Mesoangioblasts’ Efficiency in Restoring Dystrophin in a Duchenne Muscular Dystrophy Model. Stem Cells Transl Med 5: 1676–1683.

48. Quarta M, Cromie M, Chacon R, Blonigan J, Garcia V, Akimenko I, Hamer M, Paine P, Stok M, Shrager JB, et al. (2017) Bioengineered constructs combined with exercise enhance stem cell-mediated treatment of volumetric muscle loss. Nat Commun 8: 1–17.

49. Carosio S, Barberi L, Rizzuto E, Nicoletti C, Prete Z Del, Musarò A (2013) Generation of eX vivo-vascularized Muscle Engineered Tissue (X-MET). Sci Rep 3: 1–9.

50. Aguilar CA, Greising SM, Watts A, Goldman SM, Peragallo C, Zook C, Larouche J, Corona BT (2018) Multiscale analysis of a regenerative therapy for treatment of volumetric muscle loss injury. Cell Death Discov 4:.

51. Greising SM, Rivera JC, Goldman SM, Watts A, Aguilar CA, Corona BT (2017) Unwavering Pathobiology of Volumetric Muscle Loss Injury. Sci Rep 7: 1–14.

